# Deciphering the Population Ecology of Blossom Midge, *Contarinia maculipennis* Felt: An Emerging Threat to Tuberose Cultivation in India

**DOI:** 10.1101/2025.04.29.651280

**Authors:** M.C. Keerthi, Dnyaneshwar M. Firake, Y.S. Wagh, K. Ashok, K.V. Prasad, K.C. Naga, Sagar Pandit

**Affiliations:** ICAR- Indian Institute of Horticultural Research, Bengaluru, Karnataka, India; ICAR-Directorate of Floricultural Research, Pune, Maharashtra, India; ICAR-Central Potato Research Institute, Shimla, Himachal Pradesh, India; Indian Institute of Science, Education and Research, Pune, Maharashtra, India

**Keywords:** Blossom midge, life table, tuberose, soil moisture, soil types, survival rate

## Abstract

The blossom midge *Contarinia maculipennis* is a devastating pest of ornamental and vegetable crops, yet its population ecology remains poorly understood. This study employs an age-stage two-sex life table model to analyze its life history with additional focus on below-ground soil environment. Tuberose (*Agave amica*), a newly identified host, was selected for study. The average fecundity of *C. maculipennis* was 39.13 eggs, with an intrinsic rate of increase (r) of 0.137, and a mean generation time (T) of 19.73 days. The population doubling time was 5.05 days, with the highest survival rate observed in the first instar larvae (0.95), and the lowest in the adult stage (< 0.38). The population doubling time was 5.05 days. Survival was highest in first-instar larvae (0.95) and lowest in adults (< 0.38). Life expectancy peaked at age zero (e0 =15.57 days) and decreased with pivotal age. The reproductive value (Vx) peaked between 13 and 19 days post-field appearance. Pupal development was fastest, with a significantly higher adult-emergence recorded in sandy clay loam soil (9.67 days); however pupal development and adult-emergence were severely impacted in black cotton and marshy soils. Moderate soil moisture (20%) favored pupation and eclosion rate, while dry (10%) and saturated (30%) soils hindered pupation and emergence. Population projections over 90 days, starting with 133 eggs, estimated an adult population of 3,80738. These findings highlight pupal and adult stages as critical targets for management. Soil type and moisture strongly influence population dynamics, with strategies such as maintaining dry conditions between rows, deep ploughing to bury pupae, and mass trapping recommended for effective control. These insights offer practical strategies for the sustainable management of *C. maculipennis*.

## 1. Introduction

The blossom midge, *Contarinia maculipennis* Felt (Diptera: Cecidomyiidae), recently emerged as a serious pest of numerous vegetable and ornamental crops like orchids, jasmine, hibiscus, plumeria, and tuberose.^1,2^ Initially, Felt ^3^ discovered it from specimens of infested *Hibiscus* flower buds obtained from Hawaii, USA. Its entry into Hawaii is thought to have been through imported orchids from Thailand. Similarly, it spread over the world via the international trade of vegetables and cut flowers.^4^ In India, it was first reported in jasmine grown in Andhra Pradesh^5^ and Tamil Nadu^6^. It infests a wide range of plants like tomato, eggplant, pepper, and bitter melon, which shows its alarming adaptability and resilience.^7^ Female midges deposit eggs inside unopened flower buds, and the larvae develop on internal tissues, leading to gall development, deformities, and early bud drop. The impacts result in serious economic losses by lowering the aesthetic and market value of infested crops. ^2,8,9^.

Over the past few years, *C. maculipennis* infestations have resulted in serious economic damage to several high-value flower crops, posing difficulties to Indian farmers, especially in orchids^10,11^, jasmine^9,12^, and tuberose^2^. Despite its widespread occurrence, fundamental data on the population dynamics of *C. maculipennis* and the factors influencing its development remain scarce. This knowledge gap significantly hinders the development of effective management strategies in commercial crops.^11^ The recent shift of *C. maculipennis* to tuberose (*Agave amica*) as a new host, coupled with its severe outbreaks in tuberose fields across India, further emphasizes the need for targeted management strategies.^2^ Its rapid host range expansion is alarming, so it has been designated as a serious quarantine pest. Therefore, understanding its life cycle and population dynamics is essential for developing effective pest management strategies against it.

Life cycle and population dynamics can be efficiently studied by evaluating demographic traits like development rates, reproductive potentials, and survivorship using the life table.^13^ Development of sustainable management approaches often banks on identifying the most vulnerable life stages of such pests.^14^ The *C. maculipennis* pupae remain in the soil for 10–12 days, which is more than half of this insect’s lifetime. Adults with short lifespans of just 2-3 days also hide in soil cracks and crevices to protect themselves from the intense sunlight.^2^ Thus, soil conditions can be critical for this insect. Notably, the role of soil conditions has not been researched yet. Soil type and moisture content are known to substantially influence the of soil-dwelling insect’s development.^15,16^ Indeed, the research on congeneric midge species, *Contarinia nasturtii*, indicated that soil properties can be critical for its emergence behavior and survival.^17–19^ This work examines the population ecology of *C. maculipennis* using an age-stage, two-sex life table approach, while also considering the influence of below-ground soil environmental factors. Given its recent adoption of tuberose as a new host and the severe outbreaks in tuberose fields across India, we selected tuberose as the host plant to study the population ecology of *C. maculipennis*.

## 2. Materials and Methods

The life history and demography of *C. maculipennis* were studied on tuberose (var. Bidhan Ujwal) under controlled laboratory conditions (75±5 % RH; 25±1 ^0^C temperature) to understand its life cycle and development. The experiments were conducted at ICAR Directorate of Floricultural Research, Pune (India) and the experimental procedure was systematically designed to simulate the natural environment of the pest while allowing close observation of its behavior and developmental stages.

### 2.1. Studies on biological attributes of C. maculipennis on tuberose

Blossom midges were reared on tuberose buds following the methods adapted from Firake et al ^2^. For that, initially, the tuberose buds naturally infested with *C. maculipennis* were collected from the field. These buds were then placed in insect breeding dishes (Himedia: 100 × 40 mm) containing a 2.0 cm layer of moist soil at the base. The moist soil served as a medium for the larvae to pupate.

Adults started emerging after 10-12 days of pupation (Fig 1). Fresh adults were used to study the biological attributes of *C. maculipennis* on tuberose.

**Figure 1.**
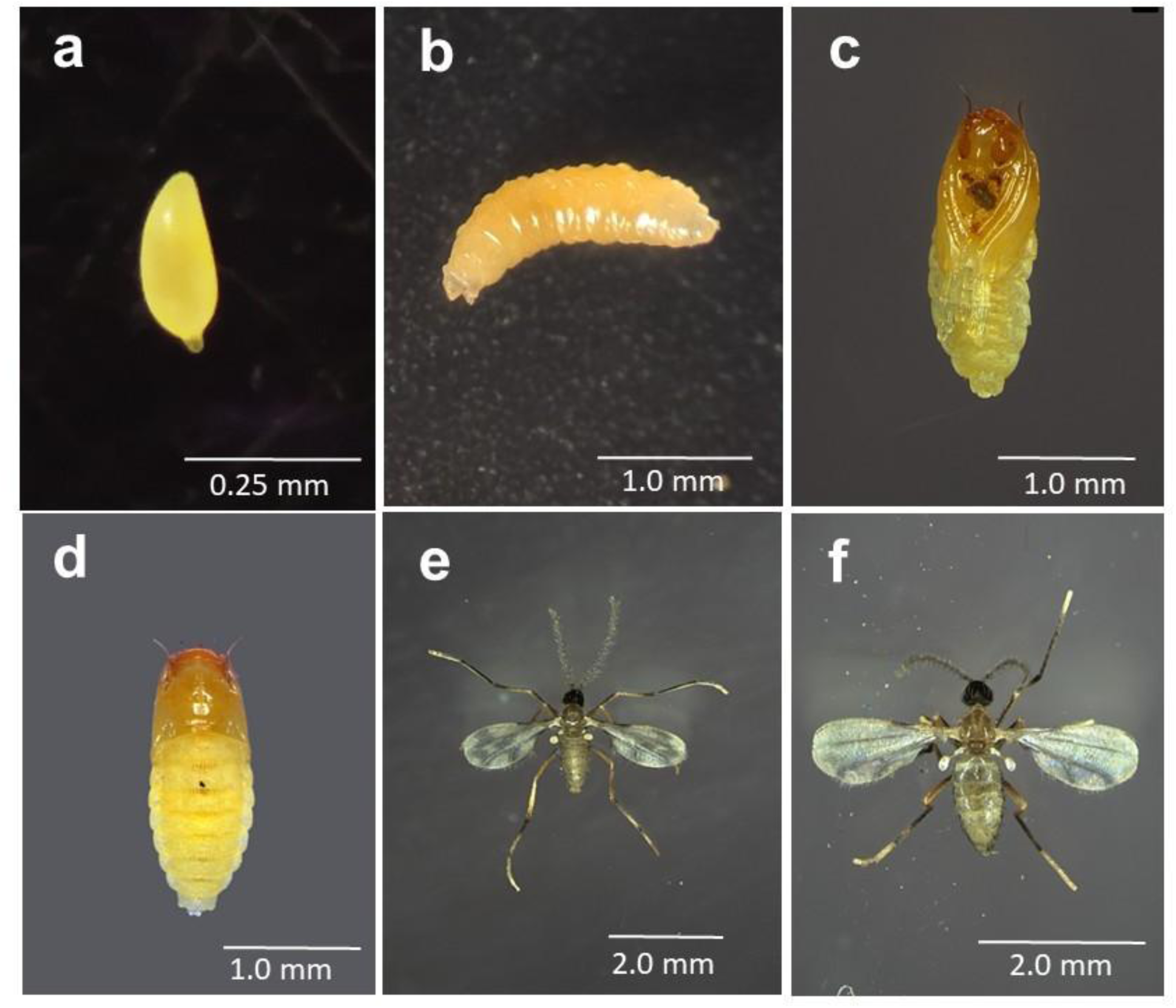
Life stages of *C. maculipennis*. **a.** Egg, **b.** Maggot. **c.** Pupa (ventral side). **d.** Pupa (dorsal side). **e.** Male, **f.** Female

Newly emerged 30 adult midges (Female: Male; 1:1 ratio) were carefully collected and released into rearing cages, which were constructed with fine netting on all sides to ensure adequate ventilation and containment. Fifty fresh tuberose spikes, each bearing newly forming buds, were introduced into the cage for oviposition inside the buds. The tuberose spikes were kept in a vase filled with a 2% sugar solution to maintain their freshness and attractiveness to the midges. To provide additional nourishment to the adult midges, a 10% honey solution (in cotton swab) was made available inside the rearing cage. This feeding setup ensured the longevity and reproductive viability of the adults.

After a 12 hour of exposure period, 15 to 20 buds from the tuberose spikes were randomly dissected under a stereomicroscope to check for the presence of eggs. Subsequently, buds were dissected every 12 hours to monitor the development of the maggots through various stages. Buds identified as infested (partially rotten or deformed) were removed from the spikes and placed in separate Himedia insect breeding dishes for pupation inside the moist soil. The breeding dishes were monitored regularly to observe the development and transition of maggots into the pupal stage. Once the midges emerged from the pupae, 20 adult flies were collected using a handheld aspirator and released into another rearing cage. Fresh tuberose spikes with newly forming buds were provided for egg-laying, following the same setup as mentioned earlier. The adult midges were fed with 10% honey solution *ad libitum*. The longevity of adult midges was recorded, with daily observations to determine the duration of their life span under laboratory conditions. The consistent monitoring of the developmental stages, from egg to adult, allowed for detailed documentation of the life cycle parameters of *C. maculipennis* on tuberose.

### 2.2. Construction of age-stage, two-sex life table of C. maculipennis

The age-stage, two-sex life table offers a significant advantage over traditional female-based life tables by eliminating many inherent inaccuracies. In this study, the life table of *C. maculipennis* was constructed using the ‘TWOSEX-MS Chart’ software,^20,21^ following the principles and methodologies outlined by Tuan et al. ^22^. This approach enabled a comprehensive analysis of population parameters, accounting for the variability among different stages and sexes. The parameters calculated included:

Age-stage-specific survival rates 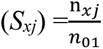;

Age-specific survival rate 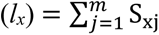;

Age-stage-specific fecundity (*f_xj_*);

Age-specific fecundity 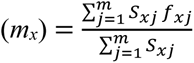

Age-specific maternity (*l_x_*\**m_x_*);

Age-stage-specific life expectancy 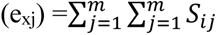;

Age-stage-specific reproductive value 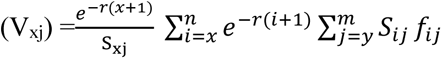;

Intrinsic rate of increase 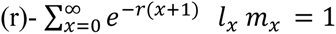;

Finite rate of increase (λ)= *e^r^*;

Net reproductive rate 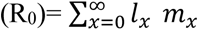;

Mean generation time 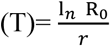.

The means, standard errors and variances of the population parameters were arrived at by bootstrapping technique (100,000 repetitions). Sigma plot 14.5 was used to create graphs.

### 2.3. Population projection

The TIMING-MSChart® program (Version 07/06/2024)^23^ was used to simulate the population growth rate and age-stage structure of *C. maculipennis* over 90 days, starting with an initial population of 133 eggs and without control.

### 2.4. Studies on the impact of soil moisture content on pupal development and the emergence of *C. maculipennis*

Since sandy clay loam is considered highly suitable for tuberose cultivation, it was chosen to investigate the effect of different soil moisture levels on the development of *C. maculipennis*. Initially, sandy clay loam soil was collected from the field and air-dried in the laboratory for one month. After drying, the soil samples were ground using a pestle and mortar and then passed through a 2-mm sieve, following the method described by Rowell^24^. The sifted soil was then placed in a hot air oven at 105 °C for 24 hours in preparation for various soil moisture treatments.^25^ After removing the dried soil samples from the oven, their weights were measured before and after saturating them with distilled water. Five soil moisture levels were established viz., 30% (Saturated), 25% (Wet), 20% (Moderate), 15% (Dry), and 10% (Very dry). The moisture content was adjusted using the following formula:

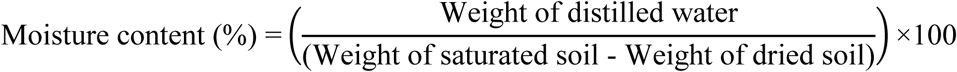

After hatching from eggs, *C. maculipennis* maggots complete their larval development within 4–5 days inside tuberose buds before exiting the buds for pupation. Thirty maggots that emerged from the buds were carefully collected from the insect culture and placed on different soil with varying moisture levels in insect breeding containers (Make: Himedia, Size: 100 × 40 mm). The larvae were allowed to burrow into the soil for pupation. The containers were weighed daily, and distilled water was added to compensate for weight loss, ensuring consistent soil moisture until the emergence of *C. maculipennis* began. Soil moisture treatments were replicated five times. Six days after the release of maggots, the containers were examined twice daily to record pupal duration and emergence rates. Adults were carefully collected using a handheld aspirator, and data collection continued until no adults emerged for seven consecutive days. At the end of the study, the containers were thoroughly inspected to count the number of pupal cases, dead larvae, or pupae (if any). Key observations, including the first day of emergence, peak emergence, and the duration of the emergence period, were recorded for each moisture level. The collected data were transformed (square and arc sine) before analysis and subjected to one-way ANOVA at a 5 and 1% significance level.

### 2.5. Studies on impact of soil types on pupal development and emergence of C. maculipennis

To assess the impact of different soil types on the biological parameters of *C. maculipennis*, six soil types were collected from a depth of 0- 20 cm. These included sandy clay loam, laterite soil, heavy black cotton soil, marshy or peaty soil, clay loam, and fine sand. Based on the observations from sandy clay loam soil, a soil moisture level of 20% was found to be the most suitable for pupal development and midge eclosion success. Therefore, a 20% moisture level was maintained for each soil type, with five replicates per treatment. The methodology for studying the biological parameters of *C. maculipennis* followed the procedure described in section 2.3. The collected data were transformed (square and arc sin) before analysis and subjected to one-way ANOVA at 5 and 1% significance level.

## 3. Results

### 3.1. Construction of age-stage, two-sex life table and population projection of C. maculipennis

Figure 1 illustrates various life stages of *C. maculipennis*. The incubation period and larval duration of *C. maculipennis* were recorded to be 1.98 ± 0.02 days and 4.54 ± 0.06 days, respectively (Table 1).

**Table 1.** Life history of blossom midge, *Contarinia maculipennis*.

| SN | Parameters | Values |
| --- | --- | --- |
| 1 | Incubation period | : 1.98±0.02 days |
| 2 | Larval period | : 4.54±0.06 days |
| 3 | Pupal duration | : 10.59±0.05 days |
| 4 | Adult pre-ovipositional period (APOP) | : 0.63±0.067 days |
| 5 | Total pre-ovipositional period from egg to first oviposition (TPOP) | : 18.31±0.108 days |
| 6 | Oviposition days | : 2.25±0.07 days |
| 7 | Adult Longevity (Male) | : 2.55±0.09 days |
| 8 | Adult Longevity (Female) | : 2.87±0.07 days |
| 9 | Fecundity | : 39.13±1.93 eggs |
| 10 | Total life cycle (Male) | : 19.26±0.15 days |
| 11 | Total life cycle (Female) | : 20.54±0.16 days |
| 12 | Intrinsic rate of increase (r) | : 0.137±0.001 No. of female offspring's /female/day |
| 13 | Finite rate of increase (lambda) | : 1.147±0.001 |
| 14 | Net reproductive rate ($R_0$) | : 15.162±0.170 No. of female offspring's /female /generation |
| 15 | Gross reproduction rate (GRR) | : 33.500±0.243 No. of female offspring's/female /generation |
| 16 | Mean generation time (T) | : 19.731±0.009 days |
| 17 | Doubling time (DT) | : 5.052±0.023 days |

The pupal stage lasted for 10.59 ± 0.05 days. The longevity of adult males was relatively shorter (2.55 ± 0.09 days), compared to females (2.87 ± 0.07 days) (Table 1). Mean female fecundity was observed to be 39.13 ± 1.93 eggs. The intrinsic rate of increase (*r*) was 0.137 ± 0.001, while the finite rate of increase (*λ*) was calculated as 1.147 ± 0.001. The net reproductive rate (R_0_) was 15.162 ± 0.170, and the gross reproduction rate (GRR) was 33.500 ± 0.243, indicating the reproductive potential of the population. The total life cycle duration, from egg to death, was 19.26 ± 0.15 days for males and 20.54 ± 0.16 days for females. The mean generation time (T) was calculated to be 19.731 ± 0.009 days, while the doubling time (DT) was 5.052 ± 0.023 days, reflecting the time required for the population to double under optimal conditions (Table 1).

The age-stage specific survival rate (S_xj_) represents the probability of individuals surviving to specific ages and developmental stages (Fig 2). The survival rates across different stages varied due to differences in developmental times among individuals. Overall, the data shows a gradual decline in survival probability from the egg stage to adulthood, reflecting cumulative mortality throughout the life stages. The highest survival rate (S_xj_) was observed during the first instar larvae (0.95), while the lowest rate was found during the adult stage (0.38 for females and 0.21 for males) followed by in pupal stage (0.62). The age-stage life expectancy (e_xj_) indicates the expected number of remaining days for individuals at each age and stage (Fig 3). Life expectancy was highest at age zero (e_0, 1_), with a value of 15.57 days, and decreased progressively with age and stage. Significant differences in life expectancy were recorded across larval instars, pupae, and adults, highlighting stage-specific mortality risks and survival potentials. The age-specific survival rate (l_x_) declined gradually throughout life, consistent with cumulative mortality trends. Female age-specific fecundity (f_x_) peaked during the mid-adult phase, which corresponds with the period of maximum reproductive output. Age-specific fecundity (m_x_) and maternity (l_x_.m_x_) exhibited similar patterns, with peak reproductive contributions occurring after the 2^nd^ days of the female’s emergence. (Fig 4). The reproductive value (V_xj_) reflects the expected future reproductive output of individuals at each age and stage. The highest reproductive value was recorded during the later adult phase of females, coinciding with the onset of egg-laying. At age zero (V_0, 1_), the reproductive value was 1.14, while on the 17^th^ day, the maximum V_xj_ value reached 34.86. Reproductive values decreased rapidly in later stages as individuals aged, indicating a decline in reproductive potential (Fig 5). Age-specific life expectancy (e_x_) started high at the egg stage and progressively declined throughout the life cycle. The reproductive value (V_x_) peaked after the 2^nd^ day of adult emergence, emphasizing the importance of this period for reproductive potential (Fig 6). This data highlights the critical role of the late adult phase in population growth and informs pest management strategies.

**Figure 2.**
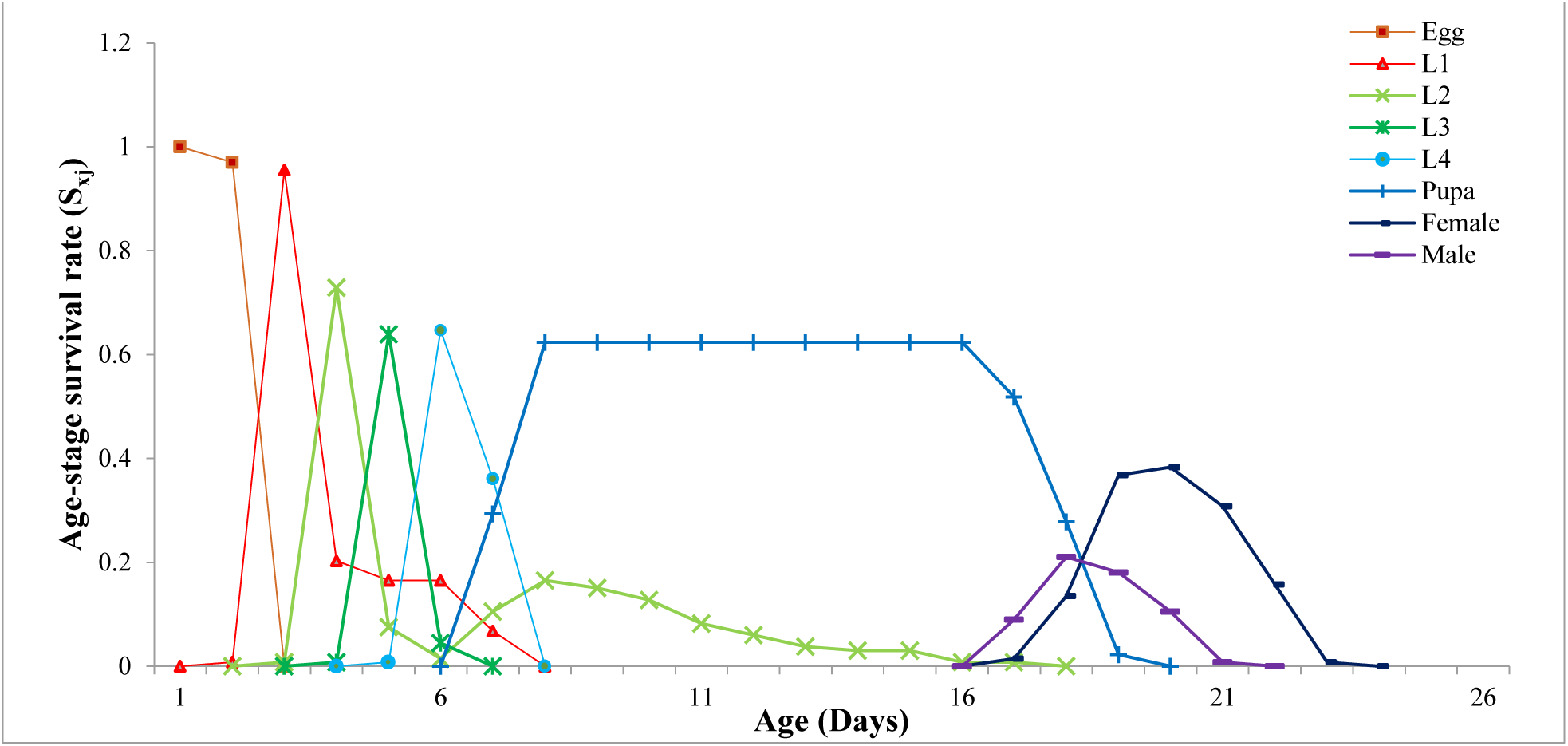
Age-stage-specific survival rate (S_xj_) of *Contarinia maculipennis* on tuberose

**Figure 3.**
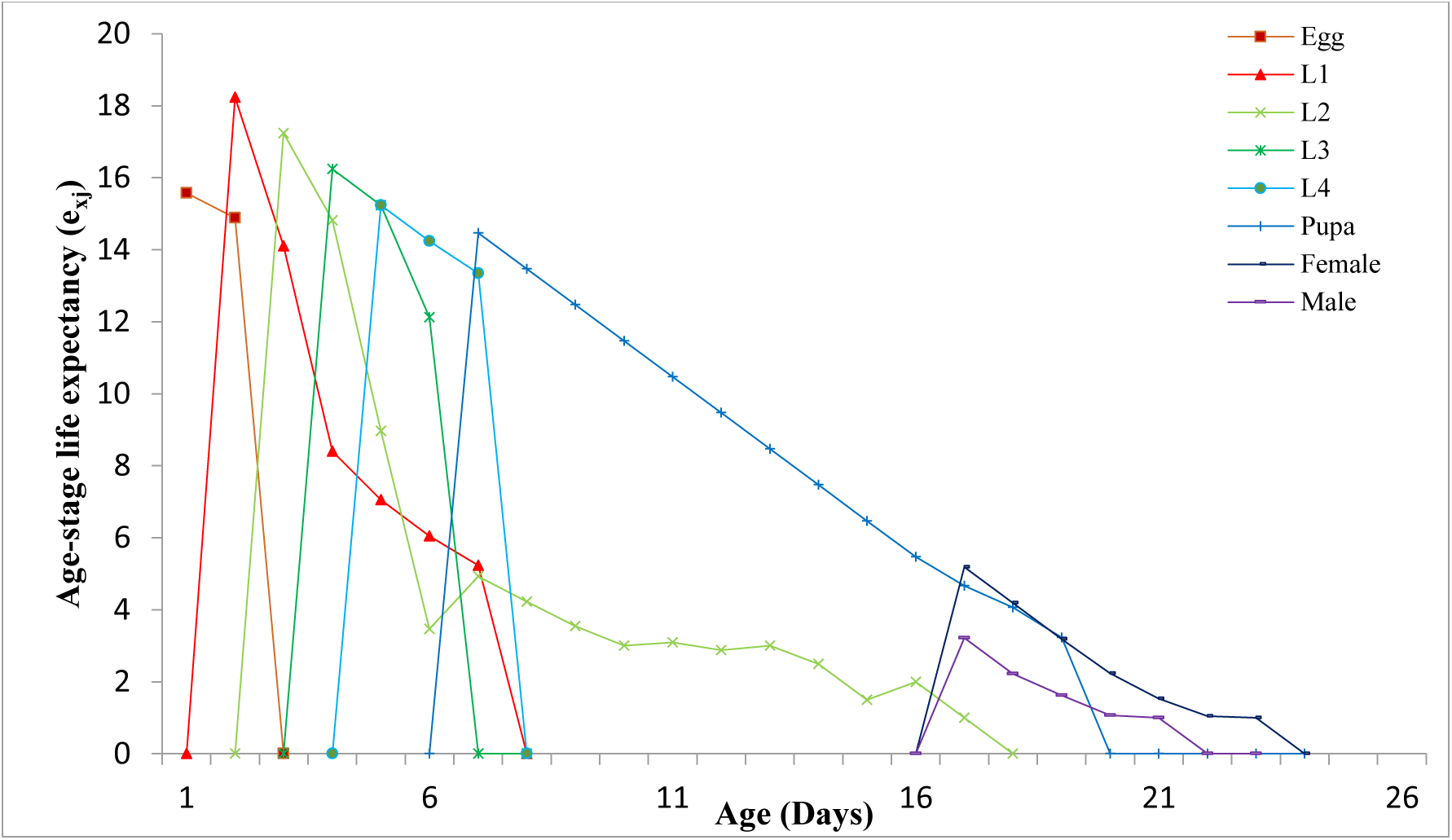
Age-stage life expectancy (e_xj_) of *Contarinia maculipennis* on tuberose

**Figure 4.**
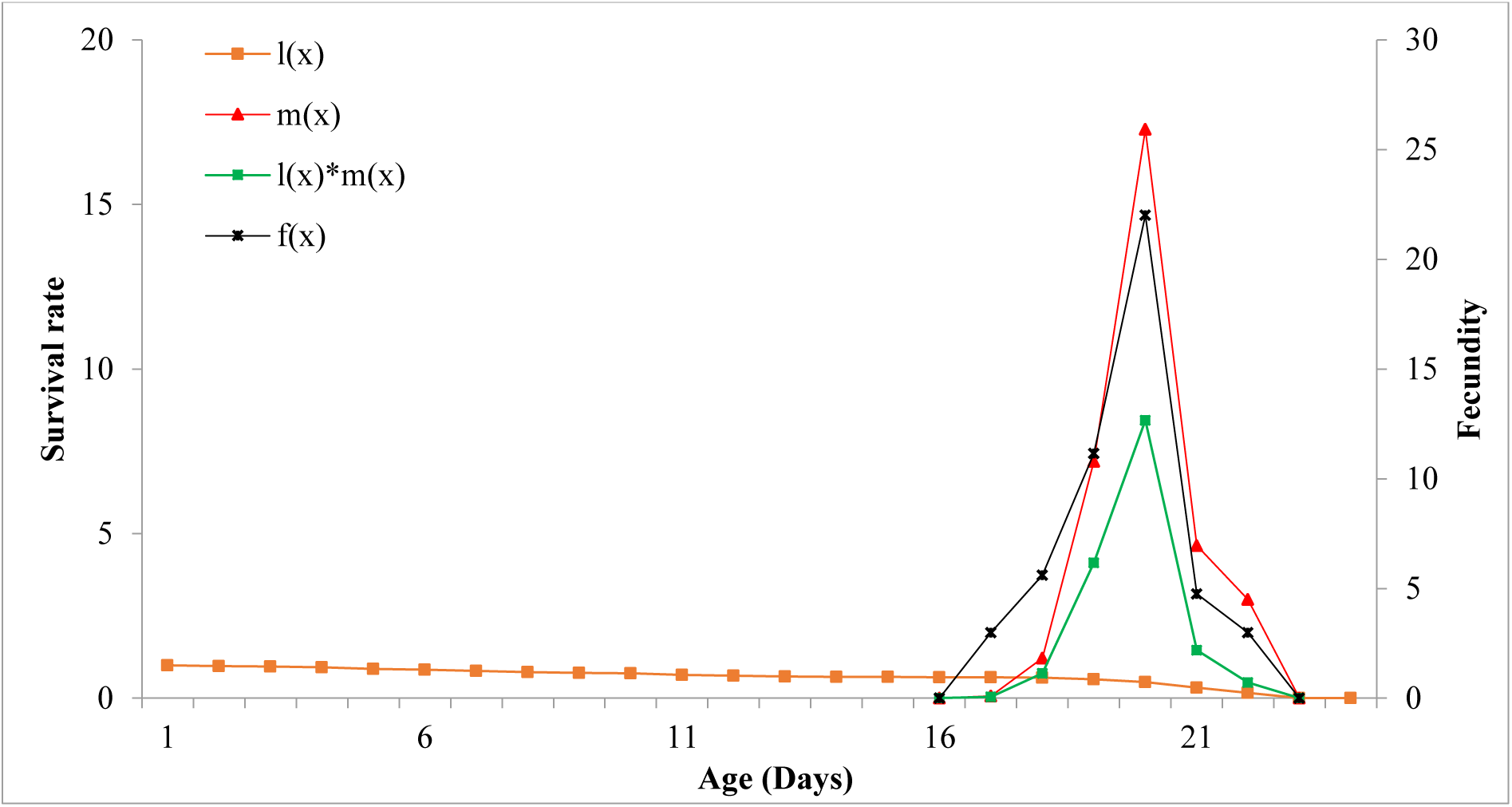
The age-specific survival rate (l_x_), age-specific fecundity (m_x_), age-specific maternity (l_x_.m_x_) and female age-specific fecundity (f_x_), of *Contarinia maculipennis* on tuberose

**Figure 5.**
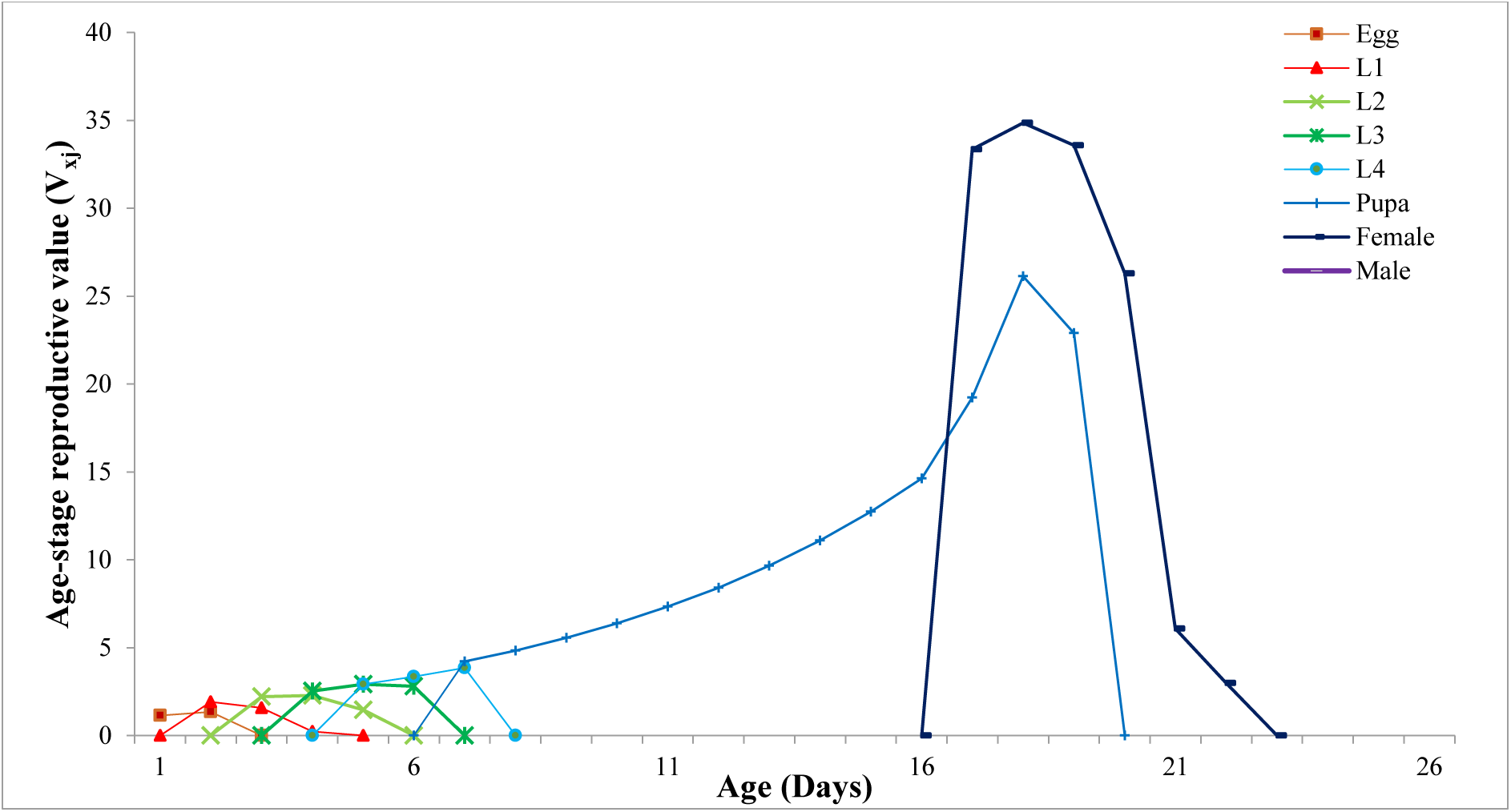
Age-stage reproductive value (V_xj_) of *Contarinia maculipennis* on tuberose

**Figure 6.**
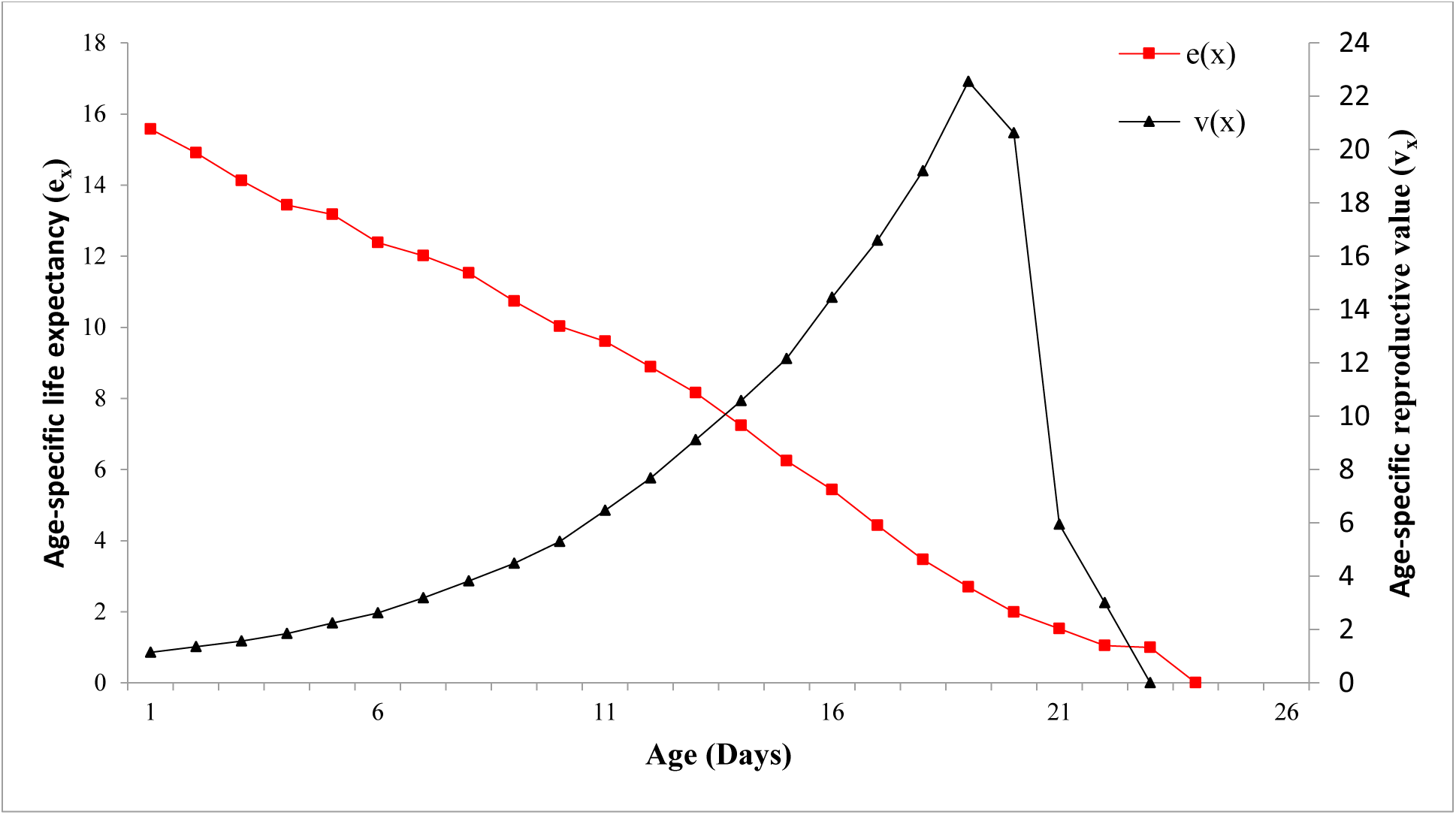
Age-specific life expectancy (e_x_) and Reproductive value (V_x_) of *Contarinia maculipennis* on tuberose

Population growth predictions for *C. maculipennis* on tuberose, generated using the TIMING- MSChart® program, are presented in Figure 7 and reveal significant variations across different life stages. Simulations indicate that adults will be ready for reproduction and egg-laying by the 16^th^ day following the first oviposition on tuberose. The predictive model suggests that if 133 eggs are present in the field, the total adult population size (N_t_) after 90 days will reach 3,80,738.

**Figure 7.**
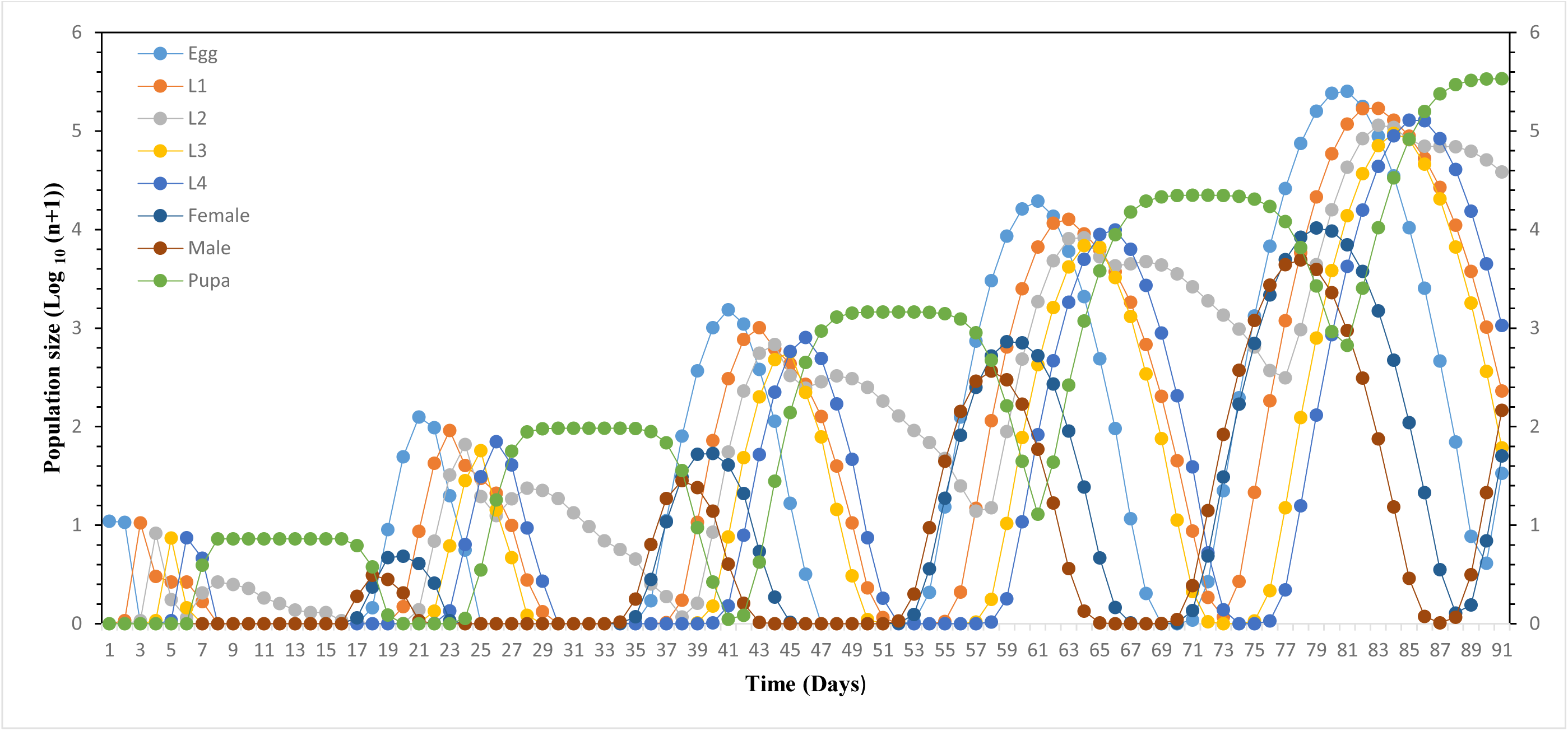
Population growth predictions of *C. maculipennis* on tuberose

### 3.2. Impact of soil moisture content on pupal development and emergence of C. maculipennis

Soil moisture levels on the emergence parameters of *C. maculipennis* revealed significant variations across all tested metrics (Table 2; p < 0.0001). The pupation success (%) was significantly higher at moderate moisture levels (20%), achieving 82.07 ± 0.78% compared to all other moisture levels. Saturated (30%) and very dry (10%) conditions resulted in the lowest pupation rates, with 14.20 ± 1.14% and 7.87 ± 0.91%, respectively. Pupal development was faster under saturated conditions (12.13 ± 0.17 days) than the other moisture levels; particularly in very dry (10% moisture) soils (11.07 ± 0.22 days). Moderate moisture yielded the highest eclosion success (88.60 ± 1.57%), while very dry soils drastically hindered emergence (19.40 ± 0.93%). The proportion of pupae near the surface was highest in moderate moisture (83.47 ± 0.62%), whereas saturated soils (64.67 ± 1.74%) showed reduced surface pupation. Adult longevity remained fairly consistent at all moisture levels, with moderate moisture producing slightly longer-lived adults (2.78 ± 0.11 days). Very dry soils yielded the shortest adult lifespan (1.62 ± 0.18 days).

**Table 2.** Impact of soil moisture content on pupal development and emergence parameters of *C. maculipennis*.

| **Moisture percentage** | **Pupation\* (%)** | **Pupal duration\*\* (days)** | **Eclosion success\* (%)** | **Proportion of pupae formation in soil depth\*\* (<0.5cm)** | **Adult longevity\*\*** |
| --- | --- | --- | --- | --- | --- |
| **30 % (Saturated)** | 14.20±1.14d<br>(22.07) | 12.13±0.17a<br>(3.48) | 25.20±1.07c<br>(30.11) | 64.67±1.74c<br>(53.55) | 2.84±0.17a<br>(1.68) |
| **25 % (Wet)** | 36.67±1.83b<br>(37.24) | 11.00±0.18b<br>(3.31) | 42.20±2.22b<br>(40.49) | 62.73±0.73c<br>(52.37) | 2.73±0.19a<br>(1.64) |
| **20 % (Moderate)** | 82.07±0.78a<br>(64.96) | 10.60±0.24b<br>(3.25) | 88.60±1.57a<br>(70.42) | 83.47±0.62a<br>(66.02) | 2.78±0.11a<br>(1.66) |
| **15 % (Dry)** | 21.07±1.22c<br>(27.28) | 10.93±0.36b<br>(3.30) | 44.60±1.86b<br>(41.89) | 72.87±1.54b<br>(58.64) | 2.20±0.14b<br>(1.48) |
| **10 % (Very Dry)** | 7.87±0.91e<br>(16.17) | 11.07±0.22b<br>(3.32) | 19.40±0.93d<br>(26.10) | 69.00±1.89b<br>(56.20) | 1.62±0.18c<br>(1.26) |
| **F Cal** | 439.62 | 5.484 | 266.80 | 38.23 | 10.83 |
| **P value** | <0.0001 | 0.004 | <0.0001 | <0.0001 | <0.0001 |

**Note:** The values in the parentheses is square transformed\*\* and arcsine transformed\*. Means followed by different letters in the same column are significantly different ( $P < 0.05$ ).

### 3.3. Impact of soil types on pupal development and emergence of C. maculipennis

The emergence parameters of *C. maculipennis* differed significantly among soil types (Table 3; p < 0.0001). Highest pupation success (82.40 ± 2.05%) was recorded in sandy clay loam soil, followed by laterite soil (73.73 ± 1.76%). In contrast, marshy soil and black cotton soils exhibited the lowest pupation rates, with 6.00 ± 0.93% and 10.80 ± 0.89%, respectively. Pupal development was fastest in sandy clay loam (9.67 ± 0.15 days), while marshy soil caused the longest pupal duration (11.87 ± 0.13 days). Fine sand and laterite soil exhibited intermediate pupal durations. Sandy clay loam yielded the highest eclosion success (87.07 ± 1.23%), whereas, the lowest eclosion success rates were observed in the black cotton soil (17.73 ± 1.10%) and in marshy soil (22.73 ± 1.04%). Highest proportion of pupae formed near the surface (up to 0.5 cm depth) in the case of black cotton soil (93.93 ± 0.64%), whereas it was lowest in the case of fine sand (11.33 ± 0.44%). Longevity was relatively consistent across most soil types, however, marshy soil and black cotton soil resulted in relatively shorter adult lifespans (2.35 ± 0.08 and 2.25 ± 0.08 days, respectively).

**Table 3.** Impact of soil types on pupal development and emergence parameters of *C. maculipennis*.

| **Moisture percentage** | **Pupation\* (%)** | **Pupal duration\*\* (days)** | **Eclosion success\* (%)** | **Proportion of pupae formation in soil depth\*\* (<0.5cm)** | **Adult longevity\*\*** |
| --- | --- | --- | --- | --- | --- |
| **Sandy clay loam** | 82.40±2.05a<br>(65.36) | 9.67±0.15d<br>(3.10) | 87.07±1.23a<br>(69.01) | 84.60±1.47c<br>(66.98) | 2.75±0.08a<br>(1.65) |
| **Laterite soil** | 73.73±1.76b<br>(59.22) | 10.27±0.13c<br>(3.20) | 77.40±1.67b<br>(61.67) | 80.73±0.55d<br>(63.97) | 2.62±0.09a<br>(1.61) |
| **Black cotton soil** | 10.80±0.89e<br>(19.12) | 11.40±0.19b<br>(3.37) | 17.73±1.10f<br>(24.86) | 93.93±0.64a<br>(75.81) | 2.25±0.08b<br>(1.49) |
| **Marshy soil** | 6.00±0.93f<br>(14.01) | 11.87±0.13a<br>(3.44) | 22.73±1.04e<br>(28.45) | 87.53±0.73b<br>(69.35) | 2.35±0.08b<br>(1.53) |
| **Clay loam** | 40.07±0.76d<br>(39.26) | 11.07±0.12b<br>(3.32) | 71.67±0.94c<br>(57.85) | 85.53±1.05bc<br>(67.70) | 2.75±0.08a<br>(1.65) |
| **Fine sand** | 63.00±0.93c<br>(52.54) | 10.00±0.15cd<br>(3.16) | 44.20±2.42d<br>(41.65) | 11.33±0.44e<br>(19.66) | 2.80±0.08a<br>(1.67) |
| **F Cal** | 419.378 | 34.35 | 340.71 | 736.43 | 8.077 |
| **P value** | <0.0001 | <0.0001 | <0.0001 | <0.0001 | <0.0001 |

*Note: The values in the parentheses are square transformed\*\* and arcsine transformed\*. Means followed by* *different letters in the same column are significantly different (P < 0.05).*

## 4. Discussion

The blossom midge, *C. maculipennis*, has a broad host range, adaptability, and the potential to spread across geographic regions through international trade, making it a significant invasive threat. Its concealed feeding habits make it difficult to assess the damage until the crop experiences sudden economic losses. Understanding the biology, reproduction, and population dynamics of any insect pest is essential for developing effective and sustainable pest management strategies.^26^ Thus, we studied detailed aspects of population ecology of *C. maculipennis* considering tuberose as a host plant. The developmental period of *C. maculipennis* on tuberose observed in this study closely aligns with previous findings on its development in jasmine (*Jasminum sambac*) and orchids, highlighting its ability to adapt to different host plants, albeit with slight variations in developmental timing. The development period of 19.26–20.54 days recorded on tuberose is longer than the 13-18 days recorded for *J. sambac*^13^, yet slightly shorter than the 21-28 days reported for *Dendrobium* orchids.^11^ The larval period (4.54 ± 0.06 days) and pupal duration (10.59 ± 0.05 days) fall within the ranges reported for other hosts, emphasizing consistent developmental patterns^27^. The female fecundity of *C. maculipennis* was observed to be 39.13 ± 1.93 eggs on tuberose. Fecundity, however, varies mainly depending on the host plant and even the same host in different locations. For example, *C. maculipennis* fecundity has been reported to be as high as 13 eggs on *Dendrobium* orchids^28^, 10 to 14 eggs on *J. sambac* at Chennai, Tamil Nadu^6^, and up to 80 eggs on *J. sambac* at Coimbatore, Tamil Nadu. These variations in fecundity may be attributed to variations in bud morphology, bud types, seasonal, host plant characteristics, and variations in measurement methods adopted by different researchers.

This is the first detailed study on demographic parameters and life table of *C. maculipennis*. The age-stage survival rate (S_xj_) was lowest at the adult stage (0.38 in females and 0.21 in males) and at the pupal stage (0.62). These findings indicate that the pupal and adult stages are the most vulnerable and critical stages to be targeted for intervention by pest management practices. The pupae were generally found to develop inside earthen cover in the soil. Thus, an increase in soil moisture may cause the pupae to cement the soil particles together, thereby preventing them from emerging as adults. Higher moisture can also increase the possibility of microbial infections in the life stages of the midges. Additionally, adult *C. maculipennis* are soft-bodied and have a short lifespan of only 2.5 to 3.0 days. Within this short period, adults must engage in several activities such as finding a mate, mating, and laying eggs in suitable habitats etc. These activities expose them to multiple challenges, which may contribute to their low survival rates. In this study, the highest fecundity (fₓ) was observed on the third-day post-emergence of females from the pupae. Thus, mass trapping of adults using sticky traps or spraying oviposition-deterrent botanicals or chemicals could significantly disrupt the midge life cycle and aid in its management. Jorgensen et al.^29^ found that yellow sticky traps effectively trapped adult *Sitodiplosis mosellana* and prevented crop damage.

The results of this study also highlight the significant impact of soil moisture levels and soil types on the pupation and emergence of *C. maculipennis*, with special reference to the interaction between soil properties and the development of midges. These findings are aligned with previous studies on other midge species and highlight several important aspects regarding soil conditions, pupal depth, and the role of moisture in emergence dynamics. Soil type and moisture together regulate midge emergence, as previously highlighted by Chen and Shelton^30^. Higher pupation success and eclosion rates in sandy clay loam and laterite soil are likely due to their balanced moisture retention properties. On the other hand, minimum pupation in black cotton and marshy soils is due to high moisture retention and poor aeration, which restricts pupal development. Additionally, moisture levels further influenced these effects, with moderate moisture (20%) producing optimal emergence rates. Both very dry and water-saturated conditions significantly decreased pupation and eclosion success, supporting reports of suboptimal emergence under extreme moisture conditions in the case of *C. nasturtii*^31^. The optimal performance of *C. maculipennis* at moderate moisture levels highlights the importance of ideal soil moisture during the pupation period. Similar patterns have been observed in wheat midges, where moderate soil moisture promotes pupal activation and adult emergence, while extreme dryness can induce dormancy or diapause^32^.

The extended pupal duration in very dry soils, as observed in this study, suggests developmental delays potentially caused by desiccation stress. Conversely, saturated soils may accelerate pupal development but at the expense of lower survival rates. The study confirmed that most pupae remained close to the soil surface (<0.5 cm) under moderate moisture conditions, supporting previous findings that shallow pupation is common in midges when soil moisture is optimum^31^. However, a higher proportion of deeper pupation (beyond 0.5 cm) was observed in black cotton soil, as its expansive nature, high plasticity, and superior moisture retention make it easier for larvae to penetrate deeper into the soil. Research indicates that burying pupae deeper than 5 cm reduces emergence rates and delays emergence time in other midge species, such as *C. nasturtii*^33^. This suggests that soil manipulation could be an effective strategy for managing *C. maculipennis* populations. Although not directly measured in this study, the results of this study indicate that high relative humidity and moderate temperatures are ideal for midge emergence, as seen in earlier research findings. For instance, Passlow^32^ observed the rapid emergence of diapause larvae at 28 °C and at 100% humidity. These conditions most likely support the physiological processes necessary for eclosion, thus the reduced emergence rates in soils with suboptimal moisture levels. The ability of *C. maculipennis* to survive in varied soil types and moisture levels is an indication of the species’ adaptability. However, extreme conditions such as prolonged dryness or excessive moisture considerably reduce survival rates, which is an indication of the need for environmental balance. Like wheat and sorghum midges, *C. maculipennis* may enter diapause in response to unfavorable conditions, ensuring long-term survival until more favorable conditions return^31,33^.

## 5. Conclusion

This study offers important insights into identifying the vulnerable stages of *C. maculipennis.* The pupae and adults were identified as the most susceptible life stages, making them crucial targets for integrated pest management strategies. Additionally, the research highlights the significant impact of soil type and moisture conditions on the development of *C. maculipennis*. Both excessive soil moisture and dryness can greatly influence midge life stages, especially during pupal development and emergence. The findings suggest that adjusting soil moisture levels during the flowering stage could serve as an effective management strategy Adopting drip irrigation systems will help maintain excessive dryness in the space between crop rows, ultimately preventing successful pupation and reducing eclosion success. Furthermore, deep ploughing or soil manipulation to bury pupae beyond their typical depth could effectively disrupt emergence cycles. Given that the adult stage is also a vulnerable target, employing mass trapping techniques with sticky traps or using oviposition-deterrent botanicals or pesticides could significantly interfere with the midge life cycle and assist in managing their population.

## Authors’ contributions

**DMF**: Conceptualization, Investigation, Methodology, Data curation, Formal Analysis, Validation, Writing original draft; **MCK**: Methodology, Data curation, Formal Analysis, Writing original draft; **YSW**: Methodology, Investigation; **AK**: Methodology, Formal Analysis; **KVP**: Funding acquisition, Project administration, Resources, Writing-review & editing; **KCN**: Data curation, Formal Analysis; **SP**: Methodology, Formal Analysis, Writing-review & editing.

## Acknowledgements

The authors would like to express their gratitude to Prof. Hsin Chi for sharing the software (TWOSEX-MS Chart pro-gram) for data analysis. ICAR funding support for the institute project “Eco-friendly Pest Management in Commercial Loose Flower Crops, Project code: 45/S/IPP/12” is duly acknowledged.

## Declarations

### Ethics approval and consent to participate

Not applicable

### Consent for publication

Not applicable

### Availability of data and material

The datasets used and/or analysed during the current study are available from the corresponding author on reasonable request

